# Data-driven models reveal the organization of diverse cognitive functions in the brain

**DOI:** 10.1101/614081

**Authors:** Tomoya Nakai, Shinji Nishimoto

## Abstract

Our daily life is realized by the complex orchestrations of diverse brain functions including perception, decision, and action. One of the central issues in cognitive neuroscience is to reveal the complete representations underlying such diverse functions. Recent studies have revealed representations of natural perceptual experiences using encoding models^1–5^. However, there has been little attempt to build a quantitative model describing the cortical organization of multiple active, cognitive processes. Here, we measured brain activity using functional MRI while subjects performed over 100 cognitive tasks, and examined cortical representations with two voxel-wise encoding models^6^. A sparse task-type encoding model revealed a hierarchical organization of cognitive tasks, their representation in cognitive space, and their mapping onto the cortex. A cognitive factor encoding model utilizing continuous intermediate features by using metadata-based inferences^7^ predicted brain activation patterns for more than 80 % of the cerebral cortex and decoded more than 95 % of tasks, even under novel task conditions. This study demonstrates the usability of quantitative models of natural cognitive processes and provides a framework for the comprehensive cortical organization of human cognition.

## Introduction

The cortical basis of daily cognitive processes has been studied using a voxel-wise encoding and decoding model approach^6^ where multivariate regression analysis is used to determine how brain activity in each voxel is modelled by target features, such as visual features^1,2^, object or scene categories^3,8,9^, sound features^5,10,11^, and linguistic information^4,12,13^. Some studies have further described the cortical (e.g. semantic) representational space that elucidates important categorical dimensions in the brain (e.g. mobile vs. nonmobile, animate vs. inanimate) and how such representations are mapped onto the cortex^3,14^. However, all previous attempts have used brain activity recorded during passive listening or viewing tasks. No study has so far been able to clarify the comprehensive cortical representations underlying active cognitive processes.

Here, we combined data-driven encoding modelling and metadata-based reverse inference to reveal such representations. Six subjects underwent functional MRI experiments to measure whole-brain blood-oxygen-level-dependent (BOLD) responses during 103 naturalistic tasks (Fig. 1a), including as many cognitive varieties as possible and ranging from simple visual detection to complex cognitive tasks such as memorization, language comprehension, and calculation (see Supp. Info for the task list and descriptions). This experimental setup aimed to extend the previous efforts at describing the semantic space^3,14^ by estimating the cognitive space that depicts the relative relationships among diverse cognitive processes. Each task was thus regarded as a sample taken from the entire cognitive space. To obtain a comprehensive representation of the cognitive space, we modelled voxel-wise responses using regularized linear regression^6^ based on two sets of features (Fig. 1b-c). First, using a task-type encoding model where tasks were represented as binary labels (Fig. 1b), we evaluated representational relationships among cognitive tasks across the cerebral cortex. Second, to further examine the generalizability of the modelling approach to any cognitive tasks, we constructed an additional cognitive factor encoding model, where each task was transformed into the 715-dimensional continuous feature space using metadata references^7^ (Fig. 1c). This allowed us to use a latent feature space for each task^6,15^ and thereby predict and decode activity for novel tasks that were not used during model training (Fig. 1d).

**Figure 1.**
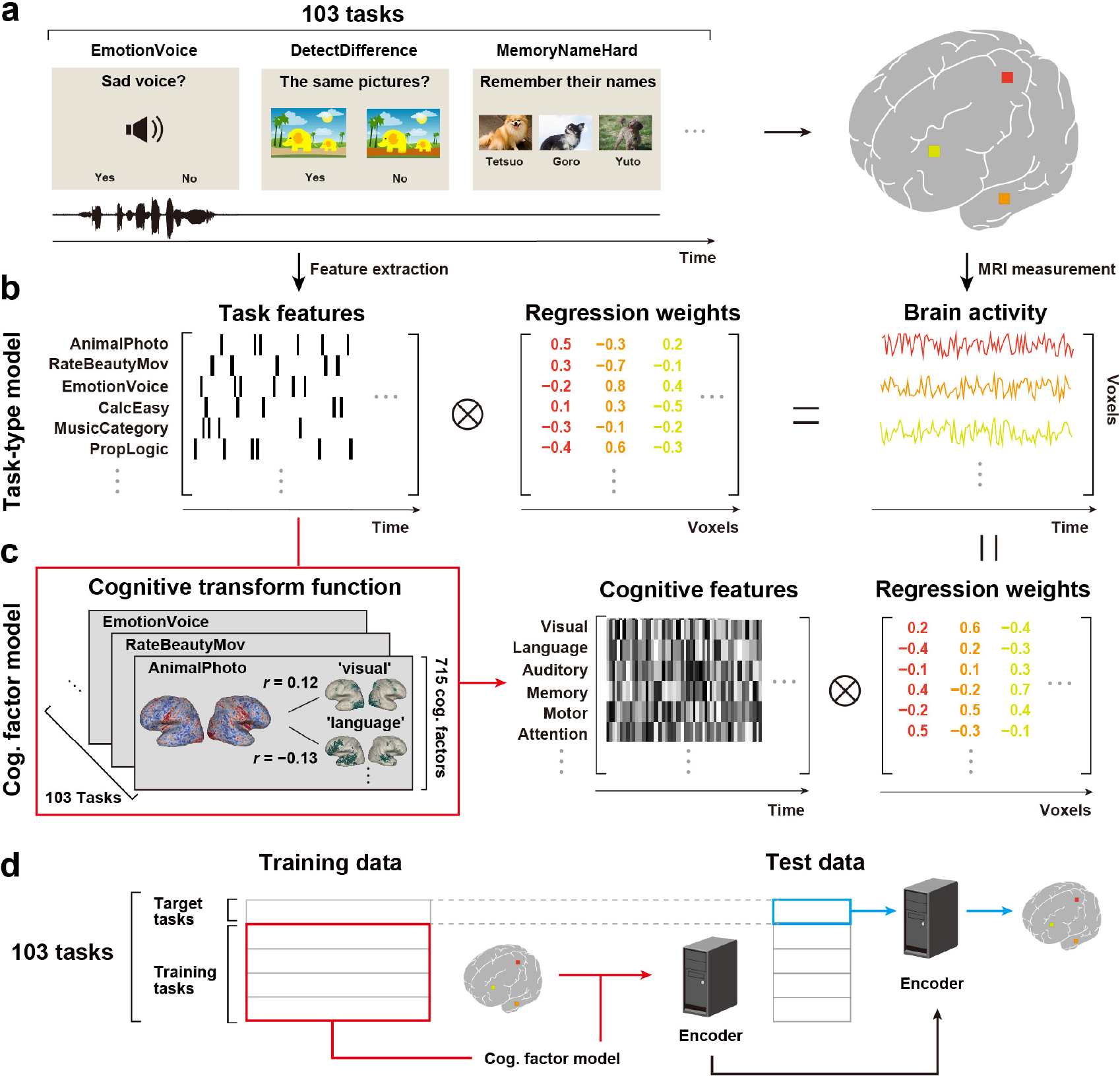
Schematic diagrams of task setting and analysis methods. **a**, Subjects performed 103 naturalistic tasks while brain activity was measured using functional MRI. **b**, Schematic of the encoding model fitting using the task-type model. **c**, Schematic of the cognitive factor model. The cognitive transform function was calculated based on correlation coefficients between the weight maps of each task and 715 metadata references^7^, and task-type features were transformed into cognitive factor features. **d**, Schematic of the encoding model fitting using the cognitive factor model for novel tasks. Target tasks were not included in the model training datasets (in red). The trained encoder provided a prediction of brain activity (in blue).

## Results

### Hierarchical organization of cognitive tasks

To examine how the cortical representations of over 100 tasks are related, we calculated a representational similarity matrix (RSM) using the estimated weights of the task-type model, concatenated across subjects (Fig. 2a). The RSM suggests that tasks form six clusters based on their representational patterns in the cerebral cortex. Task clusters were then visualized by the dendrogram obtained using hierarchical clustering analysis (HCA). The largest clusters contained tasks based on sensory modalities, such as visual (‘AnimalPhoto’, ‘MapSymbol’), auditory (‘RateNoisy’, ‘EmotionVoice’), and motor (‘PressLeft’, ‘EyeBlink’) tasks. Some clusters contained higher cognitive components, such as language (‘WordMeaning’, ‘RatePoem’), introspection (‘ImagineFuture’, ‘RecallPast’), and memory (‘MemoryLetter’, ‘RecallTaskEasy’). Tasks were further represented in sub-clusters of specific cognitive properties (Fig. 2b-d). For example, in the visual cluster, tasks with food pictures (‘RateDeliciousPic’, ‘DecideFood’) were closely located, whereas tasks with negative pictures (‘RateDisgustPic’, ‘RatePainfulPic’) formed a separate cluster, memory (Fig. 2b) tasks involving calculations (‘CalcEasy’, ‘CalcHard’) were close while those involving simple digit matching (‘MemoryDigit’, ‘MatchDigit’) formed a separate cluster, and in the introspection cluster (Fig. 2d), tasks involving imagining future and recalling past events were more closely located than tasks involving the imagination of places or faces. These results indicate hierarchically organized brain representations of cognitive tasks.

**Figure 2.**
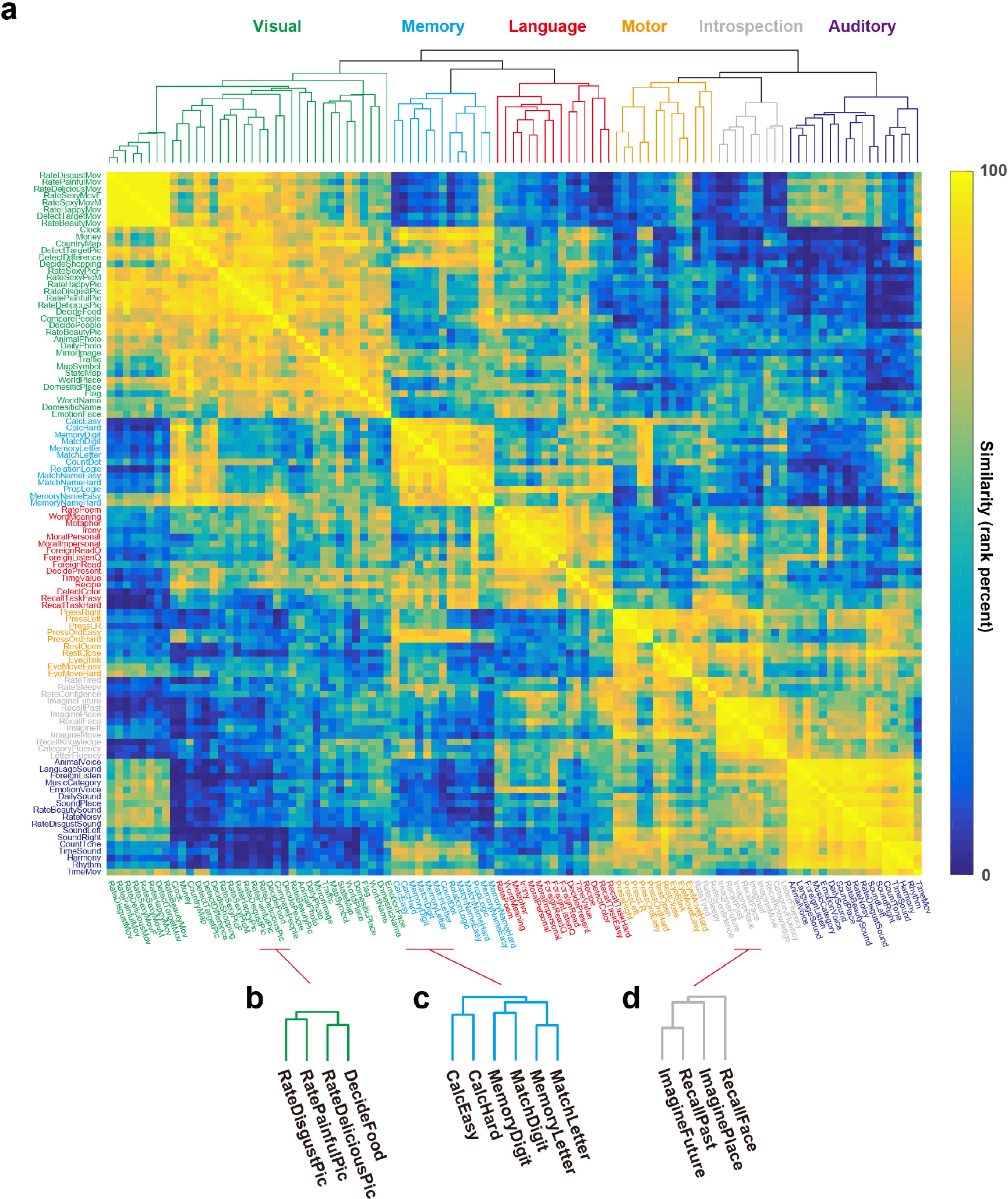
Hierarchical organization of over 100 tasks. **a**, Representational similarity matrix of the 103 tasks, reordered according to the hierarchical cluster analysis (HCA) using the task-type model weights (concatenated across subjects). The dendrogram shown at the top panel represents the results of the HCA. The six largest clusters were named after the included task types. **b-d**, Example task sub-clusters and their dendrograms in the visual (**b**), memory (**c**), and introspection clusters (**d**).

### Spatial visualization of cognitive space and its cortical mapping

The HCA reveals the relative relationships between task samples taken from the entire cognitive space. To further determine the structure and cortical organization of the cognitive space, we performed principal component analysis (PCA) with the estimated weight matrix of the task-type model, concatenated across subjects (Fig. 3 and Supplementary Fig. 1). Figure 3a shows the distributions of the tasks according to their PC coefficients, where task position is determined by the first and second PC and task colour by the first, second, and third PC (corresponding to red, green, and blue, respectively; see Fig. 3b inset). Tasks with similar representations were assigned similar colours and were closely located in the 2-dimensional space (Fig. 3a). Tasks involving movie processing are clustered on the left at the top. Tasks dedicated to image and auditory processing are located more centrally on both the left and right side, gradually shifting towards complex cognitive tasks involving language, memory, logic, and calculation at the bottom of the distribution. To further visualize cortical distributions of cognitive space representations, the voxel-wise PCs were projected to the cortical sheet of each subject (Fig. 3b and Supplementary Fig. 2, 3), using the same RGB colour scheme as in Figure 3a. For example, the occipital areas are mostly green, showing that voxels in these areas represent movie and image-related tasks (Fig. 3a). The adjacent temporal parietal junction (TPJ) tends to be coloured in red, corresponding to internal cognitive tasks involving memory and calculations. Frontal areas show intricate patterns, including language-related representations (blue) in the left lateral regions. This topographical organization was consistent across subjects (Supplementary Fig. 3), indicating that our analyses provide a broad representation of the cognitive space in the human cerebral cortex.

**Figure 3.**
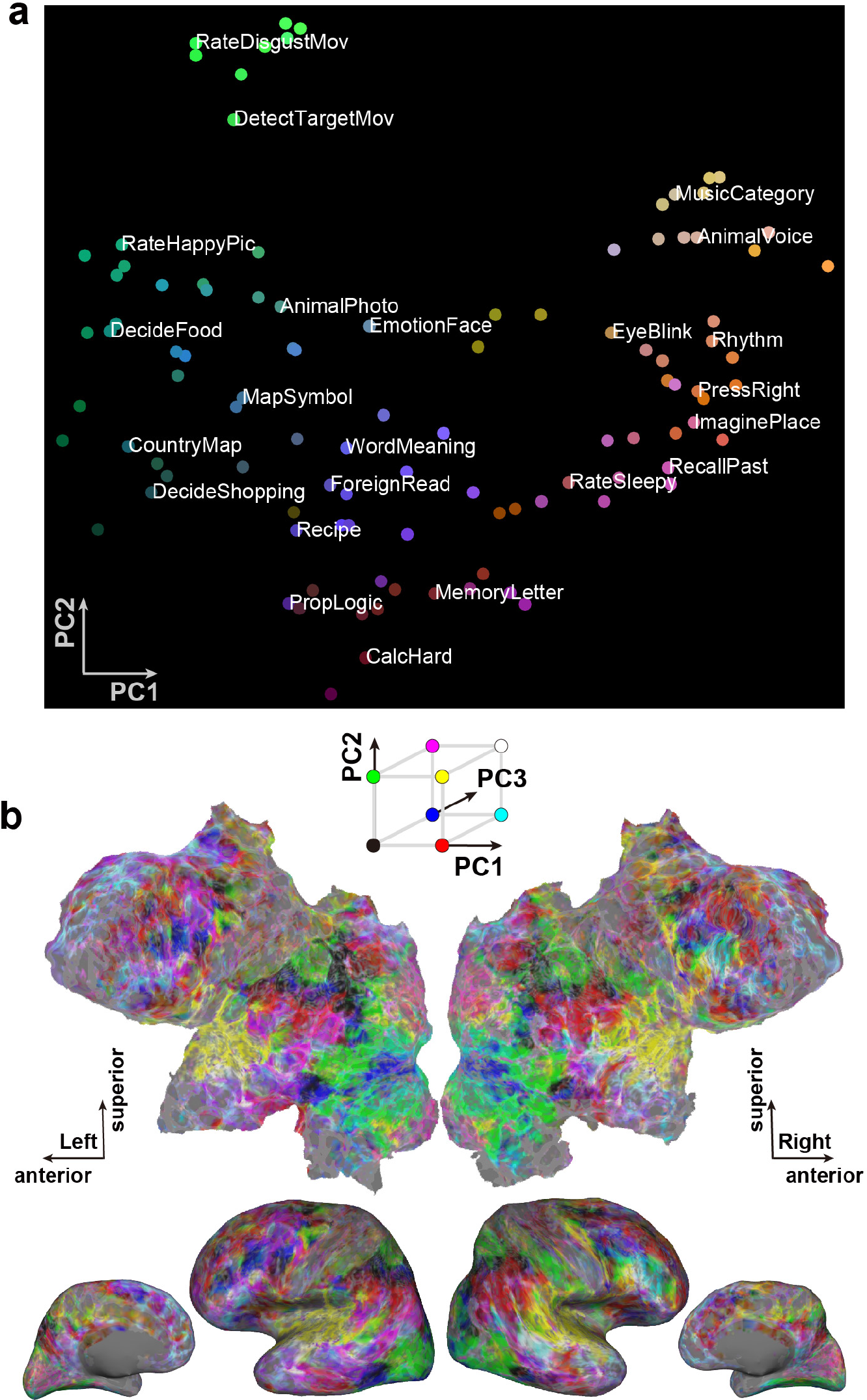
Cognitive space and cortical mapping. **a**, Colour and spatial visualization of the cognitive space. Colours indicate the loadings of the top three principal components (PC1 = auditory (red); PC2 = audiovisual (green); PC3 = language (blue)) of the task-type model weights (concatenated across subjects), mapped onto the 2-dimensional cognitive space based on the loadings of PC1 and PC2. For better visibility, only 24 tasks are labelled (in white). **b**, Cortical map of the cognitive space shown on the inflated and flattened cortical sheets of subject ID01 (Supplementary Fig. 3 shows all other subjects); PC1-PC3 are shown in red, green, and blue, respectively.

### Prediction and decoding of brain activity with the cognitive factor model

Although the task-type model can reveal distinctive relationships among tasks, it is too sparse to encompass latent and continuous features and is not generalizable to novel tasks. To tackle these issues, we transformed over 100 tasks into the 715-dimensional latent feature space using the Neurosynth database^7^ and constructed a voxel-wise cognitive factor model (Fig. 1c). To examine the generalizability of this model under novel task conditions (i.e. on a task that was not used to train the model), we trained the cognitive factor model with four fifths of the tasks (82 or 83 tasks), and predicted brain activity for the 20 or 21 remaining tasks (Fig. 1d). The model achieved significant prediction accuracy throughout the entire cortex (Fig. 4a and Supplementary Fig. 4; mean ± SD, 0.322 ± 0.042; 86.2 ± 5.1 % of voxels were significant; *p* < 0.05, FDR-corrected). To show that this cannot merely be explained by sensorimotor effects, we performed an additional encoding model analysis that regressed out visual, auditory, and motor components (see Methods). This analysis again revealed significant prediction accuracy across the cerebral cortex (mean ± SD, 0.285 ± 0.035; 82.4 ± 4.9 %; Supplementary Fig. 5), indicating that the generalizability of the cognitive factor-model stems from higher-order (i.e. not sensory) cognitive components.

**Figure 4.**
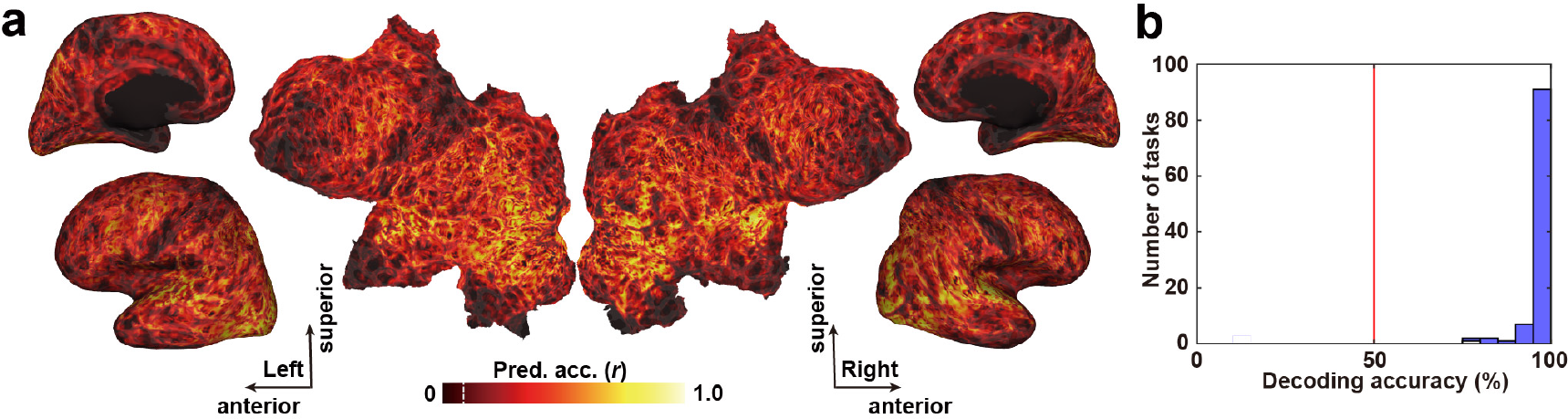
Predicting and decoding of novel tasks using the cognitive factor model. **a**, Cortical map of model prediction accuracy on inflated and flattened cortical sheets of subject ID01 (Supplementary Fig. 4 shows other subjects). Mean prediction accuracy across the cortex was 0.323 (87.2 % of voxels significant; *p* < 0.05, FDR-corrected; dashed line indicates threshold). The minimum correlation coefficient for the significance criterion was 0.0846. **b**, Histogram of task decoding accuracies for all tasks for subject ID01 (Supplementary Fig. 6 shows other subjects). The red line indicates chance-level accuracy (50 %). Blue bars show significantly decoded tasks (mean decoding accuracy, 97.5 %; 99.0 % of tasks significant; sign test, *p* < 0.05, FDR-corrected).

To further test the generalizability and task specificity of cognitive factors, we performed a task decoding analysis with novel tasks. We trained a decoding model with four fifths of the tasks and decoded the cognitive factors related to the remaining target tasks at each time point. We tested whether the decoded features were more similar to the target task than to each of the remaining 102 tasks in the cognitive space. We obtained significant decoding accuracy for novel tasks (mean ± SD, 96.5 ± 0.9 %; 98.9 ± 0.4 % of tasks were significant; sign tests, *p* < 0.05, FDR-corrected; Fig. 4b, Supplementary Fig. 6), indicating that brain activity patterns were task-specific, and that the portion of the human cognitive space our model covers is sufficient to also decode novel tasks.

## Discussion

Most previous studies using encoding or decoding model approaches have used passive viewing or listening tasks^2–4,13^, and standard neuroimaging studies using active tasks usually focus on a few conditions and examine effects of pre-assumed cognitive factors by comparing induced brain activations. While the latter strategy is a powerful way to test the plausibility of certain hypotheses, outcomes from such specialized studies could so far not elucidate the representational relationships among diverse tasks and cannot be generalized to naturalistic tasks where cognitive factors cannot be inferred in advance. Here, using over 100 naturalistic tasks that broadly sample the human cognitive space, the prediction accuracy we find for our model throughout the entire cortex is in clear contrast to the results of previous studies. While, for example, our earlier modelling attempt using a passive viewing paradigm^3^ provided significant predictions for 22 % of cortical voxels, largely restricted to the occipital and temporal areas, the cognitive factor model in the current study achieved significant predictions for about 86 % of all cortical voxels. The metadata-based inference technique used here further demonstrates the contribution of cognitive factors to these tasks^7^ and the applicability of such a data-driven approach to elucidate the brain organization of diverse cognitive functions.

While several clusters and components found in the current study have also been identified in previous multi-task studies^16–21^, we reveal a gradual shift in cognitive space, from perceptual to more complex cognitive tasks, that can only be elucidated by using our broad sampling paradigm. The subject-wise modeling also allowed the examination of the generalizability of the cognitive space, of task hierarchy, and of the representations in each subject’s brain to novel tasks; the latter may form the quantitative basis for elucidating personal traits in cognitive functions^22^. The fact that our model achieved unprecedentedly wide generalizability regarding cortical coverage and multi-task decodability indicates that our task battery represents a sufficient number of samples to probe a major proportion of the human cognitive space. Although the tasks used here do not cover the entire domain of human perception and cognition (e.g. they do not cover odour perception, speech, social interaction, etc.), our method is applicable to any arbitrary task that could be performed in a scanner, and our framework provides a powerful step forward to the complete modelling of the representations underlying human cognition.

## Methods

### Subjects

Six healthy subjects (aged 22-33 years, two females; referred to as ID01-06) with normal vision and normal hearing participated in the current experiment. Subjects were all right-handed (laterality quotient = 70-100), as assessed using the Edinburgh inventory^23^. Prior to their participation in the study, written informed consent was obtained from all subjects. This experiment was approved by the ethics and safety committee of the National Institute of Information and Communications Technology in Osaka, Japan.

### Stimuli and procedure

We prepared 103 naturalistic tasks that can be performed without any pre-experimental training (see Supplementary Data for the detailed description of each task and Supplementary Fig. 7 for the behavioural results). Tasks were selected to include as many cognitive domains as possible. Each task had 12 instances; eight instances were used in the training runs, and four instances were used in the test runs. Stimuli were presented on a projector screen inside the scanner (21.0 × 15.8 degrees of visual angle at 30 Hz). The root-mean square of auditory stimuli was normalized. During scanning, subjects wore MR-compatible ear tips. The experiment was executed in 3 days, with six runs performed on each day.

The experiment was composed of 18 runs, 12 training runs and six test runs. Each run contained 77-83 trials with a duration of 6-12 s per trial. To keep subjects attentive and engaged and to ensure all runs had the same length, a 2-s feedback for the preceding task (correct or incorrect) was presented 9-13 times per run. In addition to the task, 6 s of imaging without a task were inserted at the beginning and at the end of each run; the former was discarded in the analysis. The duration of a single run was 556 s. In the training runs, task order was pseudo-randomized, as some tasks depend on each other and were therefore presented close to each other in time (e.g. the tasks ‘MemoryDigit’ and ‘MatchDigit’). In the test runs, 103 tasks were presented four times in the same order across all six runs (but with different instances for each repetition). There was no overlap between instances in the training runs and the test runs. No explanation of tasks was given to the subjects prior to the experiment. Subjects only underwent a short training session on how to use the buttons to respond.

### MRI data acquisition

The experiment was conducted on a 3.0 T scanner (TIM Trio; Siemens, Erlangen, Germany) with a 32-channel head coil. We scanned 72 interleaved axial slices that were 2.0 mm thick, without a gap, parallel to the anterior and posterior commissure line, using a T2*-weighted gradient-echo multiband echo-planar imaging (MB-EPI) sequence^24^ [repetition time (TR) = 2000 ms, echo time (TE) = 30 ms, flip angle (FA) = 62°, field of view (FOV) = 192 × 192 mm^2^, resolution = 2 × 2 mm^2^, MB factor = 3]. We obtained 275 volumes in each run, each following three dummy images. For anatomical reference, high-resolution T1-weighted images of the whole brain were also acquired from all subjects with a magnetization-prepared rapid acquisition gradient echo sequence (MPRAGE, TR = 2530 ms, TE = 3.26 ms, FA = 9°, FOV = 256 × 256 mm^2^, voxel size = 1 × 1 × 1 mm^3^).

### fMRI data preprocessing

Motion correction in each run was performed using the statistical parametric mapping toolbox (SPM8). All volumes were aligned to the first EPI image for each subject. Low-frequency drift was removed using a median filter with a 240-s window. The response for each voxel was then normalized by subtracting the mean response and scaling it to the unit variance. We used FreeSurfer^25,26^ to identify cortical surfaces from anatomical data, and to register them to the voxels of functional data. For each subject, the voxels identified in the cerebral cortex were used in the analysis (53,345~∼66,695 voxels per subject).

### Task-type model

The task-type model was composed of one-hot vectors which were assigned 1 or 0 for each time bin, indicating whether one of the 103 tasks was performed in that period. The total number of task-type model features was thus 103.

### Encoding model fitting

In the encoding model, cortical activation in each voxel was fitted with a set of linear temporal filters that capture the slow hemodynamic response and its coupling with brain activity^2^. The feature matrix F_E_ [T × 3N] was modelled by concatenating sets of [T × N] feature matrices with three temporal delays of 2, 4, and 6 s (T = # of samples; N = # of features). The cortical response R_E_ [T × V] was then modelled by multiplying the feature matrix F with the weight matrix W_E_ [3N × V] (V = # of voxels):

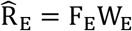

We used an L2-regularized linear regression using the training dataset to obtain the weight matrix W_E_. The training dataset consisted of 3336 samples (6672 s). The optimal regularization parameter was assessed using 10-fold cross validation, with the 18 different regularization parameters ranging from 100 to 100 × 2^17^.

The test dataset consisted of 412 samples (824 s, repeated four times). To reshape the data spanning over six test runs into the four times-repeated dataset, we discarded 6 s of the no-task period at the end of each run, as well as the 2-s feedback periods at the end of the 3^rd^ and 6^th^ test runs. Four repetitions of the test dataset were averaged to increase the signal-to-noise ratio. Prediction accuracy was calculated using Pearson’s correlation coefficient between predicted signal and measured signal in the test dataset. The statistical threshold was set at *p* < 0.05, and corrected for multiple comparisons using the false discovery rate (FDR) procedure^27^.

### Evaluation of optimal regularization parameter

To keep the scale of weight values consistent across subjects, we performed a bootstrapping procedure to assess the optimal regularization parameter used for the group HCA and PCA^4^. For each subject, we randomly divided the training dataset into training samples (80 %) and validation samples (20 %) and performed model fitting using an L2-regularized linear regression. This procedure was repeated 50 times, with the 18 different regularization parameters ranging from 100 to 100 × 2^17^. The resultant prediction accuracies were averaged across six subjects for each parameter. We selected the optimal regularization parameter which provided the highest mean prediction accuracy across subjects. This regularization parameter was used for model fitting in the group HCA and PCA.

### Hierarchical cluster analysis

For the HCA, we used the weight matrix of the task-type model concatenated across six subjects. For each subject, we selected voxels which showed a significant prediction accuracy with *p* < 0.05 (with FDR correction, 39,485~56,634 voxels per subject) and averaged three time delays for each task. RSM was then obtained by calculating Pearson’s correlation coefficients between mean brain activations of all task pairs. A dendrogram of 103 tasks was described using the task dissimilarity (1 – correlation coefficient) as a distance metric, with the minimum distance as a linkage criterion. Each cluster was labelled based on the included cognitive tasks. To obtain an objective interpretation of cluster labelling, we also performed a metadata-based inference of cluster-related cognitive factors (Supplementary Fig. 8 and Table 3).

### Principal component analysis of task-type weights

For each subject, we performed a PCA on the weight matrix of the task-type model concatenated across six subjects. We selected voxels which showed a significant prediction accuracy with *p* < 0.05 (with FDR correction, 39,485~56,634 voxels per subject) and averaged three time delays for each task. The number of meaningful PCs was determined based on the prediction accuracy with the reconstructed weight matrix (Supplementary Fig. 1). To interpret each PC, we quantified the relative contribution of each task using the PCA loadings; tasks with higher PCA loading values were regarded to contribute more to the target PC (Supplementary Table 1). Each PC was thus labelled based on these cognitive tasks. To obtain an objective interpretation of PC labelling, we also performed a metadata-based inference of PC-related cognitive factors (Supplementary Table 2). PCA loadings were also used to evaluate the representational correspondence between task clusters and PCs (Supplementary Fig. 9). To show the structure of the cognitive space, 103 tasks were mapped onto the 2-dimensional space using the loadings of PC1 (1^st^ PC) and PC2 as the x- and y-axis. The tasks were further coloured in red, green, and blue, based on the relative PCA loadings in PC1, PC2, and PC3, respectively.

To represent the cortical organization of the cognitive space for each subject, we extracted and normalized PCA scores from each subject’s voxels. The resultant cortical map indicates the relative contribution of each cortical voxel to the target PC (denoted as *PCA score map*, Supplementary Fig. 2). By combining the PCA score maps of the top three PCs of each subject, we visualized how each cortical voxel is represented by cognitive clusters. Each cortical voxel was coloured based on the relative PCA scores of PC1, PC2, and PC3, corresponding to the colour of the tasks in the 2-dimensional space.

### Cognitive factor model

To obtain a task representation using the continuous features in the human cognitive space, we transformed sparse task-type features into the latent cognitive factor feature space (Fig.1c). We used Neurosynth (http://neurosynth.org; <u>accessed 26^th^ January 2018</u>) as a metadata reference of the past neuroimaging literature^7^. From the approximately 3,000 terms in the database, we manually selected 715 terms that cover the comprehensive cognitive factors while avoiding redundancy. Specifically, we removed several plural terms which had their singular counterpart (e.g. ‘concept’ and ‘concepts’) and past tense verbs which had their present counterpart (‘judge’ and ‘judged’) in the dataset. We also excluded terms which indicated anatomical regions (e.g. ‘parietal’) (see Supplementary Data for the complete set of 715 terms). We used the reverse-inference image of the Neurosynth database for each of the selected terms. The reverse-inference image indicates the likelihood of a given term being used in a study if activation is observed at a particular voxel. Each reverse-inference image in MNI152 space was registered to the subjects’ reference EPI data using FreeSurfer^25,26^.

We calculated correlation coefficients between the weight map of each task in the task-type model and the registered reverse-inference maps. This resulted in the [103 × 715] coefficient matrix. We obtained a cognitive transform function (CTF) of each subject, by averaging the coefficient matrices of the other five subjects. The CTF is a function that transforms the feature values of 103 tasks into the 715-dimensional latent feature space. The feature matrix of the cognitive factor model was then obtained by multiplying the CTF with the feature matrix of the task-type model. Note that the CTF (and the resultant feature matrix) of each target subject was independent of their own data. The total number of cognitive factor model features was 715.

### Encoding model fitting with sensorimotor regressors

To evaluate a possible effect of low-level sensorimotor features on the model predictions, we performed an additional encoding model fitting while regressing out sensorimotor components. We concatenated motion-energy (ME) model features (visual), modulation transfer function (MTF) model features (auditory), and button response (BR) model features (motor) with the original feature matrix during the model training (see the Supplementary Methods for details). ME model features were obtained by applying 3-dimensional spatio-temporal Gabor wavelet filters to the visual stimuli^2^. MTF model features were obtained by applying spectro-temporal modulation-selective filters to the cochleogram of the auditory stimuli^28^. BR model features were obtained based on the number of button responses made by each subject. The model testing excluded the sensorimotor regressors from the concatenated feature matrix and the corresponding weight matrix. This analysis revealed that model prediction accuracy is independent of low-level sensorimotor features.

### Motion-energy model (regressor of non-interest for visual features)

The details of the ME model design have been described elsewhere^2^. First, movie frames and pictures were spatially down-sampled to 96 × 96 pixels. The RGB pixel values were then converted into the Commission International de l’Eclairage (CIE) LAB colour space, and colour information was discarded. The luminance (L*) pattern was passed through a bank of 3-dimensional spatio-temporal Gabor wavelet filters. The outputs of two filters with orthogonal phases (quadrature pairs) were squared and summed to yield local motion-energy. Motion-energy was compressed with a log-transform and temporally down-sampled to 0.5 Hz. Filters were tuned to six spatial frequencies (0, 1.5, 3.0, 6.0, 12.0, 24.0 cycles/image) and three temporal frequencies (0, 4.0, 8.0 Hz), without directional parameters. Filters were positioned on a square grid that covered the screen. The adjacent filters were separated by 3.5 standard deviations of their spatial Gaussian envelopes. The total number of ME model features was 1395.

### Modulation transfer function model (regressor of non-interest for auditory features)

A sound cochleogram was generated using a bank of 128 overlapping bandpass filters ranging from 20 to 10,000 Hz^29^. The window size was set to 25 ms, and the hop size to 10 ms. Filter output was averaged across 2 s (TR). We further extracted features from the MTF model^28^. For each cochleogram, a convolution with modulation-selective filters was calculated. The outputs of two filters with orthogonal phases (quadrature pairs) were squared and summed to yield local modulation energy^2^. Modulation energy was log-transformed, averaged across 2 s, and further averaged within each of the 10 non-overlapping frequency ranges logarithmically spaced along the frequency axis. The filter outputs of upward and downward sweep directions were used. Modulation-selective filters were tuned to five spectral modulation scales (Ω = 0.50, 1.0, 2.0, 4.0, 8.0 cyc/oct) and five temporal modulation rates (ω = 4.0, 8.0, 16.0, 32.0, 64.0 Hz). The total number of MTF model features was 1000.

### Button response model (regressor of non-interest)

The BR model was constructed based on the number of button responses within 1 s for each of the four buttons, with the right two buttons pressed by the right thumb and the left two buttons pressed by the left thumb. The total number of BR model features was four.

### Decoding model fitting

In the decoding model, the cortical response matrix R_D_ [T × 3V] was modelled by concatenating sets of [T × V] matrices with temporal delays of 2, 4, and 6 s. The feature matrix F_D_ [T × N] was modelled by multiplying the cortical response matrix R_D_ with the weight matrix W_D_ [3V × N]:

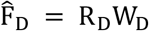

The weight matrix W_D_ was estimated using an L2-regularized linear regression with the training dataset, following the same procedure as for the encoding model fitting.

### Encoding and decoding with novel tasks

In order to examine the generalizability of our models, we performed encoding and decoding analyses with novel tasks which were not used in the model training (Fig. 1d). We randomly divided the 103 tasks into five task groups. A single task group contained 20-21 tasks. We performed five independent model fittings, each with a different task group as the target. From the training dataset, we excluded the time points during which the target tasks were performed, and those within 6 s after the presentation of the target tasks. In the test dataset, we used only the time points during which the target tasks were performed, and those within 6 s after the presentation of the target tasks. This setting allowed us to assume that the activations induced by the target task group and those induced by the other four task groups (training task groups) did not overlap, and it enabled us to investigate the prediction and decoding accuracy for the novel tasks. We performed the encoding and decoding model fitting with the training task groups composed of 82-83 tasks. For the model testing, we concatenated the predicted responses or decoded features of the five task groups. Responses or features for the time points that were duplicated were averaged across the five task groups. Note that encoding and decoding with novel tasks was only possible with the cognitive factor model, because the original tasks needed to be transformed into the latent feature space.

For the decoding analysis with novel tasks, we measured the similarity between the CTF of each task and each decoded cognitive factor vector using Pearson’s correlation coefficient for each time point. We refer to the correlation coefficient as the *task score*^12^. We then calculated the time-averaged task scores for each task using the one-vs.-one method. For each target task, a series of binary classification was performed between the target task and each of the remaining 102 tasks. Decoding accuracy was then calculated as a percentage that the target task had a higher task score in this procedure. Statistical significance of decoding accuracy was tested for each task using the sign test (*p* < 0.05, with FDR correction).

### Code and data availability

The MATLAB code used in the current study and the datasets generated during and/or analysed during the current study are available from the corresponding author upon reasonable request.

## Acknowledgments

We thank MEXT/JSPS KAKENHI (grant numbers 17K13083 and JP18H05091 in #4903 (Evolinguistics) for T.N., and JP15H05311 for S.N.) as well as JST CREST JPMJCR18A5 and ERATO JPMJER1801 (for S.N.) for the partial financial support of this study. The funders had no role in the study design, data collection and analysis, decision to publish, or preparation of the manuscript.

## Author contribution

T.N. and S.N. designed the study; T.N. collected and analysed the data; T.N. and S.N. wrote the manuscript.

## Author Information

The authors declared no competing interests. Correspondence and requests for materials should be addressed to S.N

## Supplementary information and Extended data

**Supplementary Figure 1.**
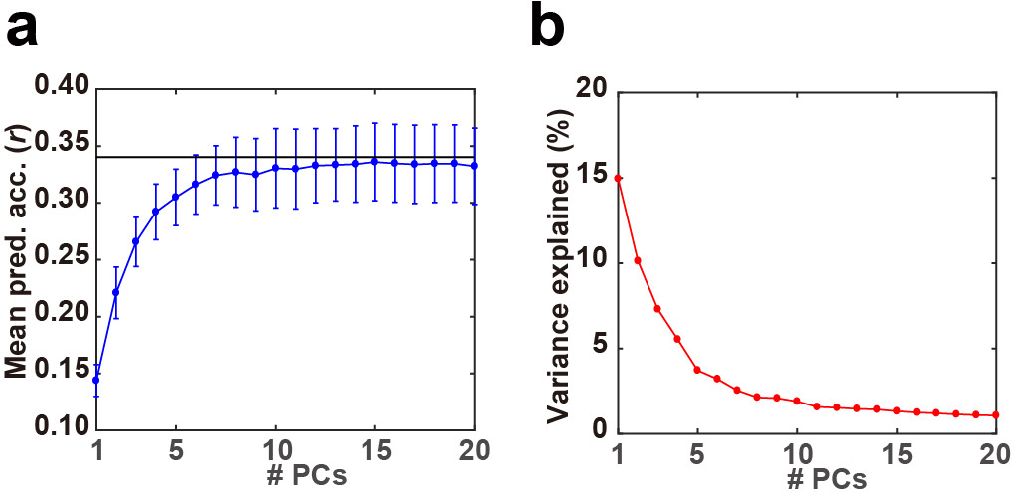
Determination of the number of meaningful components. **a**, Prediction accuracy of the task-type model with reconstructed weight matrices using principal component analysis (PCA) results, averaged across six subjects. The black line indicates the original prediction accuracy averaged across six subjects. Error bars indicate SD. **b**, Variance explained in the PCA. The explained variance of the original weight matrix of the task-type model was plotted for each PC. Note that the explained variance is common for all subjects in the group PCA.

### Determination of the number of meaningful components

To determine the number of meaningful principal components (PCs), we calculated the prediction accuracy of the task-type model with the reconstructed weight matrix, using a restricted number of PCs. For each subject, the weight matrix W_EK_ [N × V] was reconstructed by multiplying the 1st to Kth component of the PCA loading matrix V_K_ [N × K] with the PCA score matrix U_K_ [V × K]:

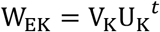

The cortical response R_E_ [T × V] was then modelled by multiplying the feature matrix F_E_ [T × N] with the reconstructed weight matrix W_EK_:

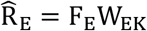

Note that the feature matrix in this analysis did not include the three temporal delay components, since the original weight matrix used in the PCA was averaged across delays. The prediction accuracy was calculated using Pearson’s correlation coefficient between predicted signals and measured signals in the test dataset. For comparison, we also calculated the prediction accuracy with the original weight matrix.

This analysis showed that the prediction accuracy reached a plateau with the top eight PCs (Supplementary Fig. 1a), at a total explained variance of 49.5 % (Supplementary Fig. 1b). We thus analysed the dominant cognitive components based on these eight PCs.

**Supplementary Figure 2.**
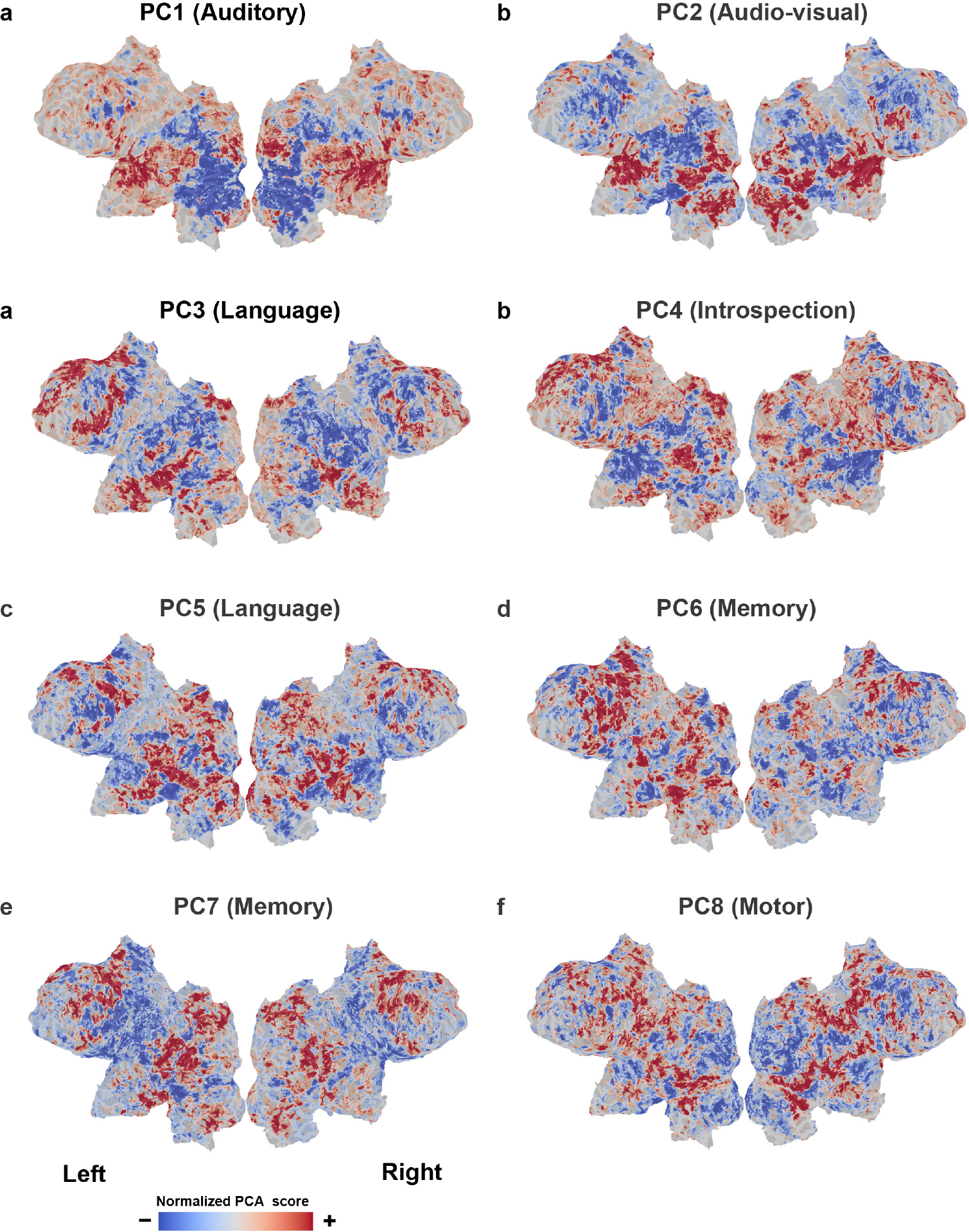
PCA score maps. Normalized principal component analysis (PCA) scores of PC1-8 projected onto the flattened cortical sheets of subject ID01.

### Cortical representation of principal components

To assess the cortical regions related to the top PCs, we projected normalized PCA scores onto the cortical maps (Supplementary Fig. 2). PC1 (auditory component, according to Supplementary Table 1) had large weights in the superior temporal regions. PC2 (audiovisual component) had large weights in the superior temporal and occipital regions. PC3 and PC5 (language components) had large weights in the frontal and inferior temporal regions. PC4 (introspection component) had large weights in the medial frontal and cingulate regions. PC6 and PC7 (memory components) had large weights in the frontal and parietal regions. PC8 (motor component) had large weights in the pericentral regions. These PCA score maps were further used to evaluate the cognitive factors related to each PC.

**Supplementary Figure 3.**
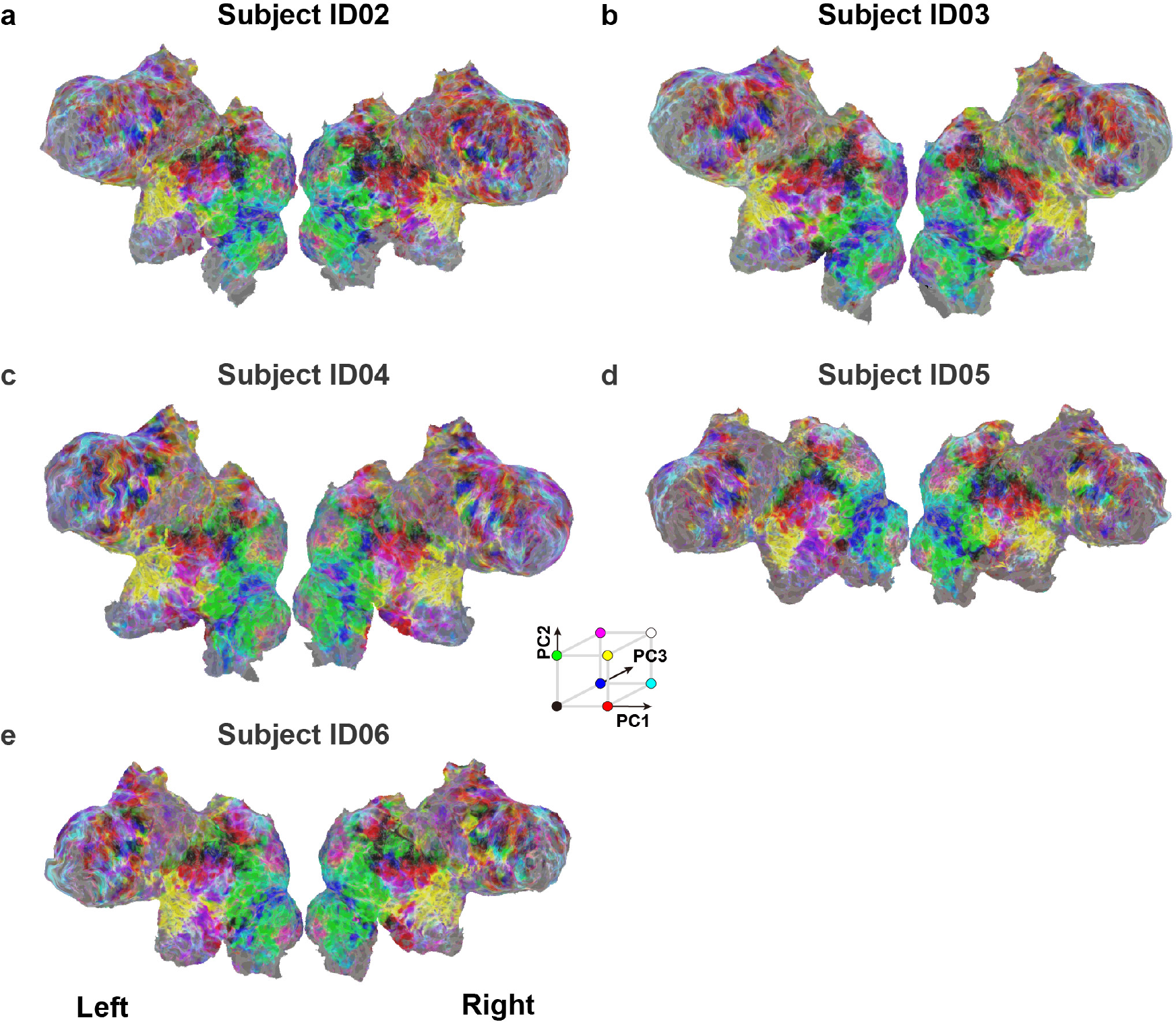
Cortical mapping of the cognitive space. Cortical maps of the cognitive space are shown on the inflated and flattened cortical sheets of subjects ID02-ID06, visualized using scores of the top three principal components (PC1-PC3) in red, green, and blue, respectively.

**Supplementary Table 1.** Top tasks related to each principal component

|  | Top 10 tasks |
| --- | --- |
| PC1<br>(auditory) | 'SoundLeft', 'ForeignListen', 'SoundRight', 'CountTone', 'RestClose', 'Harmony', 'RateBeautySound', 'ImagineMove', 'Rhythm', 'RateNoisy' |
| PC2<br>(audiovisual) | 'RateSexyMovM', 'RatePainfulMov', 'RateHappyMov', 'RateDisgustMov', 'RateSexyMovF', 'RateDeliciousMov', 'RateBeautyMov', 'DetectTargetMov', 'RateBeautySound', 'RateNoisy' |
| PC3<br>(language) | 'MoralPersonal', 'Irony', 'MoralImpersonal', 'ForeignListenQ', 'WordMeaning', 'ForeignReadQ', 'Metaphor', 'DecidePresent', 'RatePoem', 'DomesticName' |
| PC4<br>(introspection) | 'ImagineMove', 'ImaginePlace', 'ImagineIf', 'RecallPast', 'RestOpen', 'PressLeft', 'PressRight', 'EyeBlink', 'PressLR', 'RecallFace' |
| PC5<br>(language) | 'MoralPersonal', 'MoralImpersonal', 'EyeMoveHard', 'EyeMoveEasy', 'Metaphor', 'Irony', 'ForeignRead', 'EyeBlink', 'PropLogic', 'ForeignReadQ' |
| PC6<br>(memory) | 'RecallKnowledge', 'LetterFluency', 'ImagineIf', 'CategoryFluency', 'ForeignRead', 'ImaginePlace', 'MemoryNameEasy', 'MemoryNameHard', 'MemoryLetter', 'MemoryDigit' |
| PC7<br>(memory) | 'RecallPast', 'RecallTaskEasy', 'RecallTaskHard', 'ImagineFuture', 'MatchNameHard', 'MatchNameEasy', 'DomesticPlace', 'DetectTargetMov', 'ImaginePlace', 'TimeMov' |
| PC8<br>(motor) | 'ButtonOrdEasy', 'ButtonOrdHard', 'MatchNameHard', 'MatchNameEasy', 'DomesticName', 'WorldName', 'PressRight', 'RateSexyPicM', 'ForeignListenQ', 'RateSexyPicF' |

Top 10 tasks with the largest PCA loadings for PC1 to PC8.

**Supplementary Table 2.** Top cognitive factors related to each principal component

|  | Top 10 cognitive factors in the NeuroSynth database |
| --- | --- |
| PC1<br>(auditory) | 'auditory', 'sound', 'listening', 'speech', 'acoustic', 'pitch', 'audiovisual', 'speech perception', 'auditory visual', 'music' |
| PC2<br>(audiovisual) | 'auditory', 'pitch', 'sound', 'acoustic', 'listening', 'audiovisual', 'musical', 'music', 'hearing', 'multisensory' |
| PC3<br>(language) | 'comprehension', 'semantic', 'sentence', 'language', 'linguistic', 'syntactic', 'word', 'language comprehension', 'reading', 'lexical' |
| PC4<br>(introspection) | 'default', 'default mode', 'mode', 'mode network', 'dmn', 'network dmn', 'default network', 'motor', 'somatosensory', 'sensorimotor' |
| PC5<br>(language) | 'theory mind', 'mind tom', 'tom', 'mentalizing', 'mind', 'mode', 'default mode', 'default', 'mode network', 'visual' |
| PC6<br>(memory) | 'eye', 'task', 'production', 'phonological', 'working memory', 'saccade', 'preparation', 'language', 'load', 'speech production' |
| PC7<br>(memory) | 'memory', 'retrieval', 'default', 'episodic', 'default mode', 'mode', 'memory retrieval', 'autobiographical memory', 'navigation', 'mode network' |
| PC8<br>(motor) | 'hand', 'finger', 'action', 'movement', 'finger movements', 'motor', 'motor task', 'action observation', 'sensorimotor', 'viewing' |

Top 10 cognitive factors in the Neurosynth database for PC1 to PC8, based on the correlation coefficients between each PC score map and the 715 registered reverse-inference maps.

### Interpretation of cognitive factors related to principal components

To interpret the plausible cognitive factors related to the target sub-clusters, we used Neurosynth (http://neurosynth.org; <u>accessed 26^th^ January 2018</u>) as a metadata reference of the past neuroimaging literature^7^. Each reverse-inference image in the Neurosynth database in MNI152 space was registered to the subjects’ reference EPI data using FreeSurfer^25,26^. For each PCA score map obtained with the task-type model, we calculated Pearson’s correlation coefficients between the PCA score map and the 715 registered reverse-inference maps, resulting in a cognitive factor vector with 715 elements. Terms with higher correlation coefficient values were regarded to contribute more to the target PC.

Several top PCs distinguished a class of sensorimotor components to the others, such as auditory (PC1; tasks with high loading, e.g. ‘SoundLeft’, ‘Harmony’), audiovisual (PC2; ‘RatePainfulMov’, ‘RateHappyMov’), and motor components (PC8; ‘ButtonOrdEasy’, ‘PressRight’), while the other PCs distinguished higher-order cognitive components such as language (PC3; ‘Irony’, ‘WordMeaning’; PC5; ‘MoralPersonal’, ‘Metaphor’), memory (PC6; ‘RecallKnowledge’, ‘MemoryNameEasy’; PC7; ‘RecallPast’, ‘RecallTaskHard’), and introspection components (PC4; ‘ImagineMove’, ‘ImaginePlace’) (Supplementary Table 1).

To obtain an objective interpretation of the estimated PCs, we performed a metadata-based reverse inference of the cognitive factors related to each PC using the NeuroSynth database^7^. The top 10 terms for each PC provided an interpretation of the relevant cognitive components, with PC1 (auditory) showing a high correlation with auditory-related terms (e.g. ‘auditory’, ‘sound’), PC2 (audiovisual) showing a high correlation with auditory- and vision-related terms (e.g. ‘auditory’, ‘audiovisual’), PC3 (language) showing a high correlation with language-related terms (‘language’, ‘sentence’), and PC8 (motor) showing a high correlation with motor-related terms (‘movement’, ‘motor’) (Supplementary Table 2). These results were largely consistent with the interpretations based on the high loading task types.

For the other components, the interpretation based on NeuroSynth provided a more detailed description. Although we labelled PC4 as ‘introspection component’ based on the high loading task types, PC4 showed a high correlation with default mode-related terms in the Neurosynth database (‘default mode’, ‘dmn’). Moreover, PC5 was labelled as ‘language component’ but showed a high correlation with theory of mind-related terms (‘theory mind’, ‘mentalizing’). PC6 and PC7 were both labelled as ‘memory components’, but the former was more related to the working memory component (‘language’, ‘working memory’) and the latter was related to the default mode-related component (‘memory’, ‘default’). These results indicate that the metadata-based reverse inference reliably captured fundamental cognitive factors involved in the diverse cognitive tasks.

**Supplementary Figure 4.**
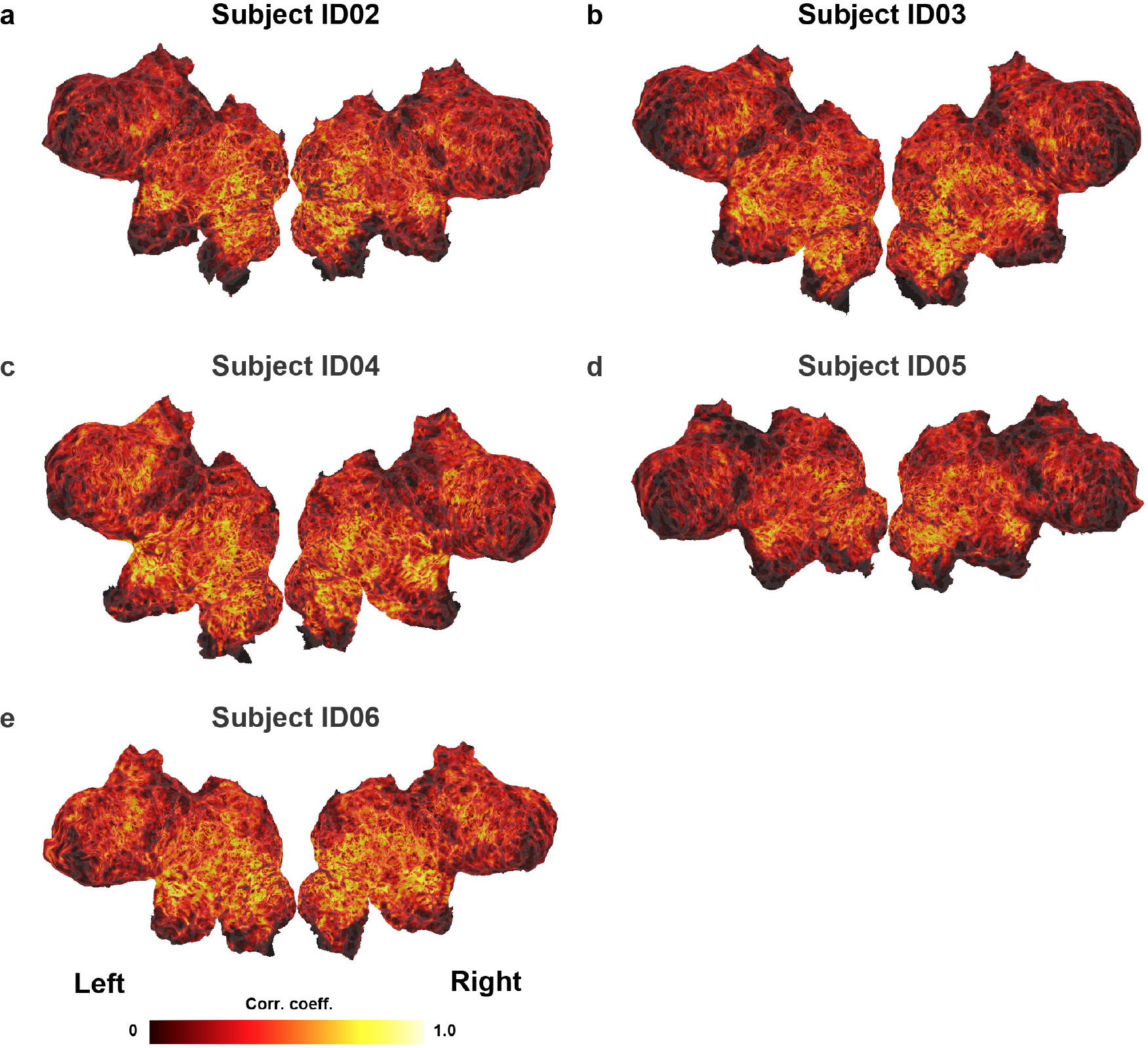
Prediction accuracy using the cognitive factor model under novel task conditions. The cortical map is shown on the flattened cortical sheets of subjects ID02-ID06 (mean prediction accuracy and percent significant voxels; ID02, 0.311 and 86.4 %; ID03, 0.316 and 84.5 %; ID04, 0.357 and 90.7 %; ID05, 0.252 and 77.2 %; ID06, 0.373 and 91.4 %; *p* < 0.05, FDR-corrected, the minimum correlation coefficient for the significance criterion ranged from 0.0833 to 0.0872 for each individual subject).

**Supplementary Figure 5.**
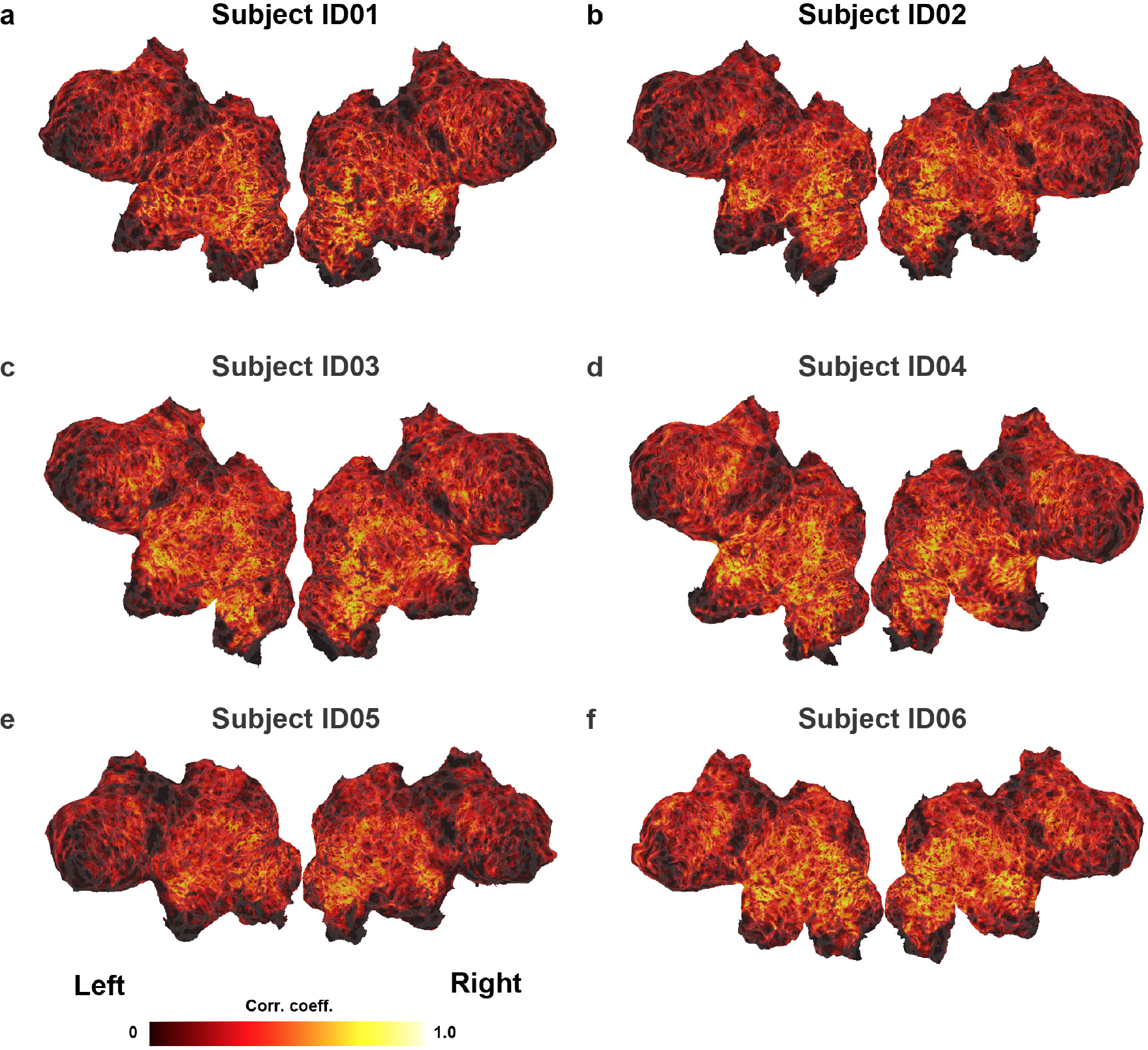
Prediction accuracy of the cognitive factor model excluding sensorimotor features. The cortical map is shown on the flattened cortical sheets of subjects ID01-ID06 (mean prediction accuracy and percent significant voxels; ID01, 0.285, 82.3 %; ID02, 0.273, 82.0 %; ID03, 0.283, 81.7 %; ID04, 0.315, 86.9 %; ID05, 0.226, 73.9 %; ID06, 0.327, 87.6 %; *p* < 0.05, FDR-corrected, the minimum correlation coefficient for the significance criterion ranged from 0.0844 to 0.0881 for each individual subject).

**Supplementary Figure 6.**
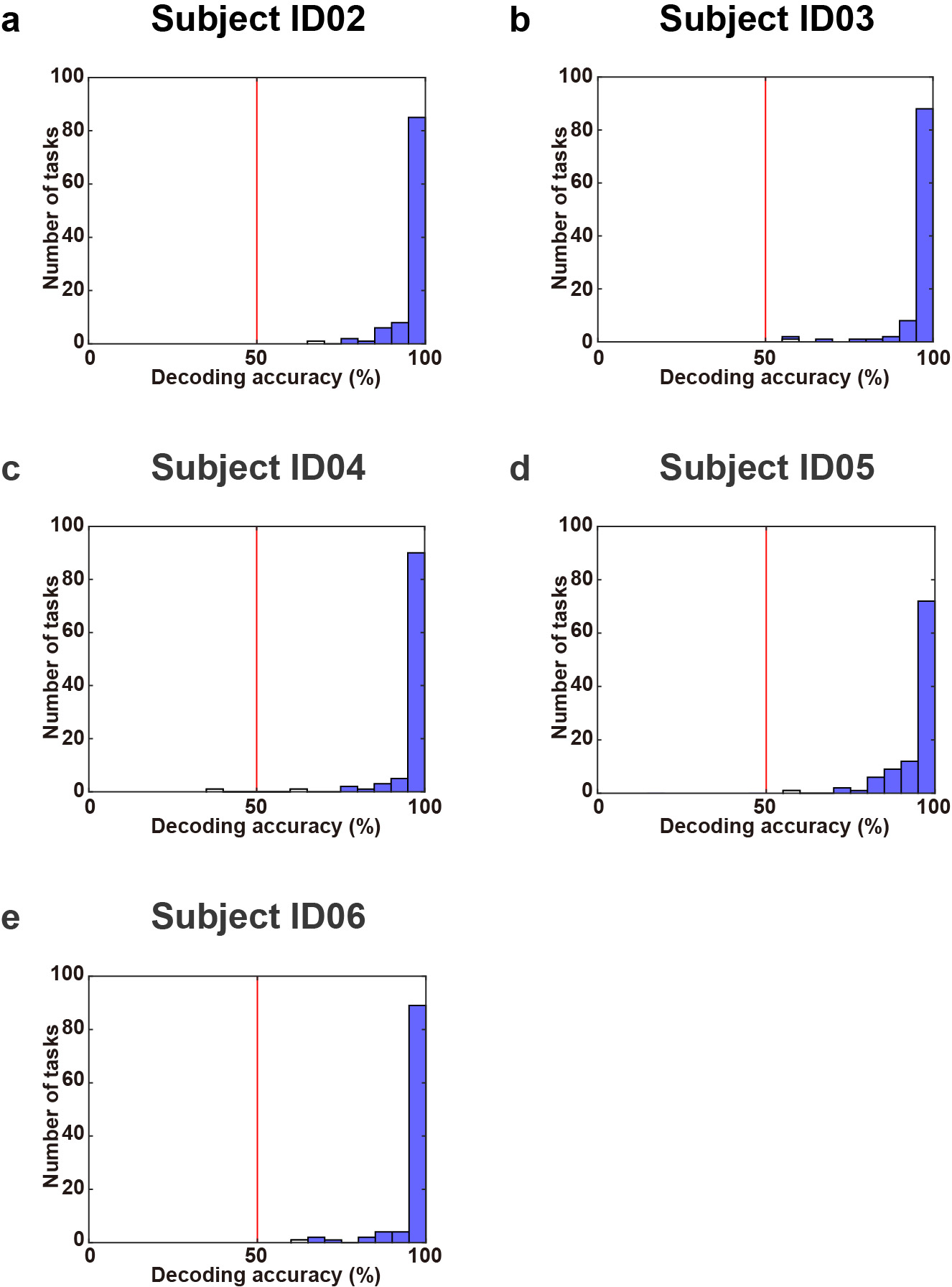
Decoding accuracy of novel tasks. Histogram of decoding accuracies of over 100 tasks obtained using the cognitive factor model with novel tasks, for subjects ID02-ID06. The red line indicates chance-level accuracy (50 %). Bars showing significantly decoded tasks are filled in blue (mean decoding accuracy and percent significant tasks; ID02, 96.7 %, 99.0 %; ID03, 96.5 %, 99.0 %; ID04, 96.7 %, 98.0 %; ID05, 94.8 %, 99.0 %; ID06, 96.6 %, 99.0 %; sign tests, *p* < 0.05, FDR-corrected).

**Supplementary Figure 7.**
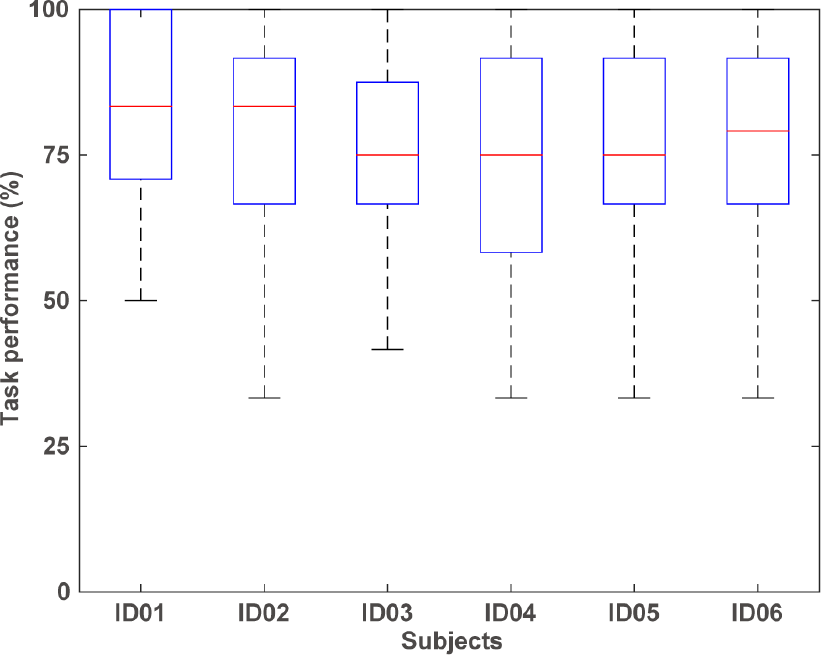
Behavioural results. Box plots of accuracy for 48/103 tasks are shown for subjects ID01-06 Each box shows the median (red), the interquartile range (blue), and the maximum and minimum values.

### Behavioural results

To show that the tasks used in the current study were sufficiently natural and easy to perform, we analysed the behavioural performance for 48/103 tasks (Supplementary Fig. 7). The 48 tasks were selected because only these tasks presented a single ‘yes or no’ question. All subjects performed these tasks significantly better than chance level (mean ± SD, 77.9 ± 2.8 %; Wilcoxon signed-rank tests, *p* < 0.05, FDR-corrected), indicating that they understood the tasks without any pre-experimental training or explanation. We also confirmed that the subjects did not have any difficulty in understanding the task settings via self-reports after the experiment.

**Supplementary Figure 8.**
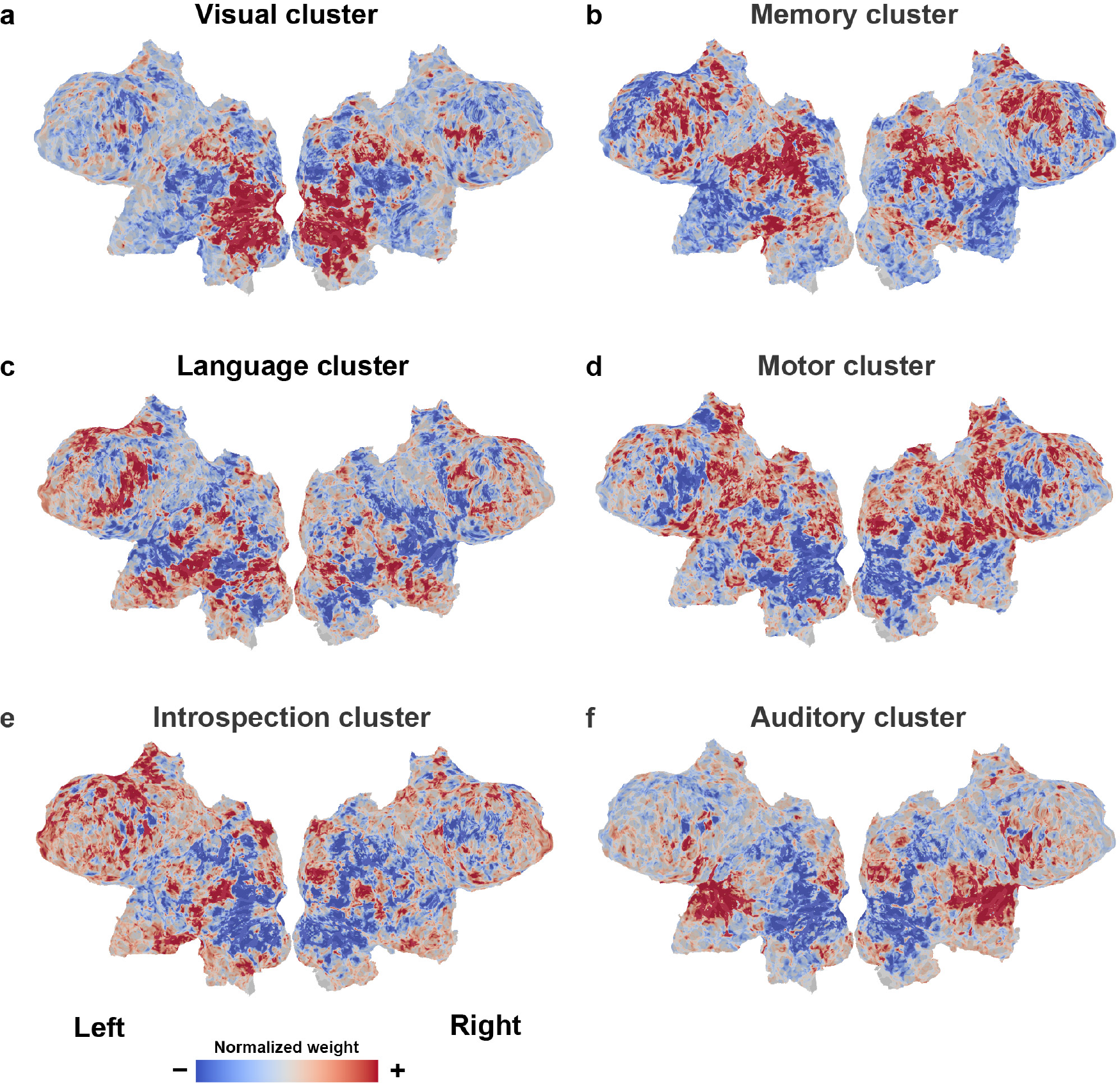
Cortical maps of task clusters. Normalized cortical maps of weight matrices using the task-type model, showing the visual (**a**), memory (**b**), language (**c**), motor (**d**), introspection (**e**), and auditory (**f**) clusters.

### Cortical representation of task clusters

To assess the brain regions related to each task cluster, we examined the weight matrix for only those tasks that are within each of the six largest task clusters (Supplementary Fig. 8). The weight values of the target clusters were averaged across tasks and normalized across voxels. This indicates the relative contribution of each cortical voxel to the target task cluster. The task cluster related to visual processing showed large weights in occipital regions, the cluster related to auditory processing showed large weights in the superior temporal regions, and the one related to memory processing showed large weights in the frontal and parietal regions. The language cluster showed large weights in the left frontal and inferior temporal regions, the motor cluster showed large weights in the pericentral regions, and the task cluster that was related to introspective processes showed large weights in the medial frontal and cingulate regions. These cluster weight maps were further used to evaluate the cognitive factors related to each task cluster.

**Supplementary Table 3.** Top cognitive factors related to each task cluster

|  | Top 10 cognitive factors in the NeuroSynth database |
| --- | --- |
| Visual cluster | ‘visual’, ‘object’, ‘face’, ‘motion’, ‘viewing’, ‘perceptual’, ‘vision’, ‘visual motion’, ‘early visual’, ‘visual stream’ |
| Memory cluster | ‘working memory’, ‘task’, ‘calculation’, ‘load’, ‘attentional’, ‘memory wm’, ‘numerical’, ‘spatial’, ‘arithmetic’, ‘memory load’ |
| Language cluster | ‘reading’, ‘language’, ‘comprehension’, ‘sentence’, ‘semantic’, ‘word’, ‘linguistic’, ‘native’, ‘syntactic’, ‘lexical’ |
| Motor cluster | ‘finger’, ‘motor’, ‘hand’, ‘sensorimotor’, ‘somatosensory’, ‘movement’, ‘motor imagery’, ‘execution’, ‘finger movements’, ‘motor task’ |
| Introspection cluster | ‘default’, ‘default mode’, ‘mode’, ‘mode network’, ‘autobiographical’, ‘dmn’, ‘network dmn’, ‘default network’, ‘self’, ‘autobiographical memory’ |
| Auditory cluster | ‘auditory’, ‘sound’, ‘pitch’, ‘listening’, ‘acoustic’, ‘speech’, ‘music’, ‘musical’, ‘audiovisual’, ‘speech perception’ |

Top 10 cognitive factors in the Neurosynth database for each of the six task clusters, based on the correlation coefficients between the task weight map and the 715 registered reverse-inference maps.

### Top cognitive factors related to each task cluster

We labelled each task cluster of the HCA (e.g. ‘visual cluster’ or ‘language cluster’) based on the included task types. To avoid arbitrariness, we performed a metadata-based objective evaluation of the task clusters using the NeuroSynth metadata^7^. For each of the cortical maps of the task cluster weight matrix, we calculated Pearson’s correlation coefficients with the 715 registered reverse-inference maps, resulting in a cognitive factor vector with 715 elements. Terms with higher correlation coefficient values were regarded as contributing more to the target cluster.

The top 10 terms for most task clusters were consistent with our interpretation based on the included task types (Supplementary Table 3). The visual cluster showed a high correlation with vision-related terms (e.g. ‘visual’, ‘perception’), the memory cluster with working memory-related terms (‘working memory’, ‘memory’), and the language cluster showed a high correlation with language-related terms (‘language’, ‘reading’). The motor cluster showed a high correlation with motor-related terms (‘movement’, ‘motor’), and the auditory task cluster showed a high correlation with auditory-related terms (‘auditory’, ‘listening’). The introspection cluster showed a high correlation with default mode-related terms (‘default mode’, ‘default’), in line with our interpretation of the introspection PC (PC4). These results suggest that data-driven reverse inference is effective in providing an objective evaluation of the cognitive factors underlying different task clusters.

**Supplementary Figure 9.**
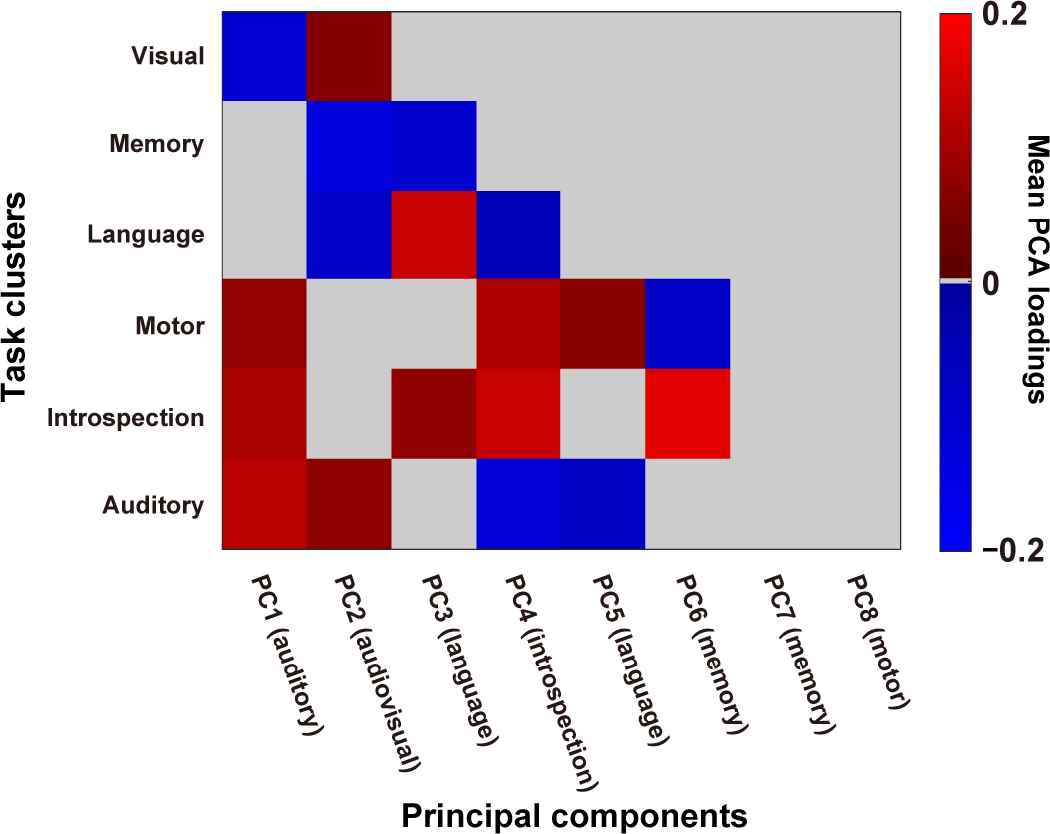
Relationship between principal components and task clusters. Mean PCA loadings of the tasks in the six largest clusters, plotted for each of the top eight PCs. Task clusters with positive and negative PCA loadings are shown in red and blue, respectively (significantly different from zero; sign tests, *p* < 0.05, FDR-corrected). Task clusters with non-significant PCA loadings are shown in grey.

### Correspondence between task clusters and principal components

To examine the relationship between the HCA and PCA results, we calculated the relative contribution of the top eight PCs to the six largest task clusters (Supplementary Fig. 9). We averaged the PCA loadings of the tasks included in each of the target clusters. We found that the top four PCs corresponded well to the related clusters (with mean PCA loadings significantly larger than zero; sign test, *p* < 0.05, FDR-corrected). PC1 (auditory component) contributed to the auditory cluster; PC2 (audiovisual component) contributed to the auditory and visual clusters, PC3 (language component) contributed to the language cluster; and PC4 (introspection component) contributed to the motor and introspection clusters. These results indicate the representational correspondence between the HCA and PCA results.

### Description of each task

1. PressRight Subjects pressed the buttons (with their right hand) as many times as possible. Duration: 8 s.
2. PressLeft Subjects pressed the buttons (with their left hand) as many times as possible. Duration: 8 s.
3. PressLR Subjects pressed the buttons (with their right or left hand) as many times as possible. Duration: 8 s.
4. RestOpen Subjects did not perform any task, with their eyes open. Duration: 10 s.
5. RestClose Subjects did not perform any task, with their eyes closed. Duration: 10 s.
6. EyeBlink Subjects blinked their eyes as many times as possible. Duration: 8 s.
7. RateTired Subjects rated how tired they were by pressing one of the four buttons. Duration: 6 s.
8. RateConfidence Subjects rated how confident they were about their accuracy on the previous task by pressing one of the four buttons. Duration: 6 s.
9. RateSleepy Subjects rated how sleepy they were by pressing one of the four buttons. Duration: 6 s.
10. ImagineFuture Subjects imagined their future situation (e.g. ‘Imagine your next weekend’). Duration: 8 s.
11. ImagineIf Subjects imagined they were some other living thing. Duration: 8 s.
12. ImagineMove Subjects imagined their body moving. Duration: 8 s.
13. ImaginePlace Subjects imagined a certain place. Duration: 8 s.
14. RecallPast Subjects recalled a past event. Duration: 8 s.
15. RecallKnowledge Subjects recalled as many names as possible which have a given property (e.g. recall as many Japanese river names as possible). Duration: 10 s.
16. RecallFace Subjects recalled the face of somebody. Duration: 8 s.
17. LetterFluency Subjects recalled as many words as possible starting with a given letter. Duration: 10 s.
18. CategoryFluency Subjects recalled as many words as possible belonging to a given word category. Duration: 10 s.
19. Clock Subjects looked at a photo of a clock, and judged whether the indicated time matched the time displayed above the photo. Duration: 6 s.
20. AnimalPhoto Subjects looked at a photo of an animal, and judged whether its name matched the name displayed above the photo. Duration: 6 s.
21. AnimalVoice Subjects listened to the voice of an animal, and judged whether its name matched the name shown on the screen. Duration: 6 s.
22. Money Subjects looked at a photo of money, and judged whether the indicated amount matched the amount displayed above the photo. Duration: 8 s.
23. Traffic Subjects looked at a photo of a traffic sign, and judged whether its meaning matched the meaning indicated above the photo. Duration: 6 s.
24. EmotionFace Subjects looked at a photo of a face with a specific emotion, and judged whether the emotion matched the emotion indicated above the photo. Duration: 6 s.
25. EmotionVoice Subjects listened to a voice with a specific emotion, and judged whether the emotion matched the emotion indicated above the photo. Duration: 6 s.
26. Flag Subjects looked at a photo of a national flag, and judged whether the country matched the country indicated above the photo. Duration: 6 s.
27. MapSymbol Subjects looked at a photo of a map symbol, and judged whether its meaning matched the meaning indicated above the photo. Duration: 6 s.
28. CalcEasy Subjects solved an easy arithmetic problem using single digits. Duration: 8 s.
29. CalcHard Subjects solved a difficult arithmetic problem using two-digit numbers. Duration: 10 s.
30. DailyPhoto Subjects looked at a photo of a tool used daily, and judged whether its name matched the name displayed above the photo. Duration: 6 s.
31. DailySound Subjects listened to the sound of tool used daily, and judged whether its name matched the name displayed above the photo. Duration: 6 s.
32. CountDot Subjects counted the number of presented dots. Duration: 8 s.
33. CountTone Subjects counted the number of presented tones. Duration: 8 s.
34. CountryMap Subjects looked at a photo of a country map and judged whether its name (nation) matched the name displayed above the photo. Duration: 6 s.
35. StateMap Subjects looked at a photo of a state (prefecture) map and judged whether its name matched the name displayed above the photo. Duration: 6 s.
36. RateSexyPicF Subjects looked at a photo of a female, and rated how sexy they thought she was. Duration: 6 s.
37. RateSexyPicM Subjects looked at a photo of a male, and rated how sexy they thought he was. Duration: 6 s.
38. RateSexyMovM Subjects viewed a movie of a male, and rated how sexy they thought he was. Duration: 10 s.
39. RateSexyMovF Subjects viewed a movie of a female, and rated how sexy they thought she was. Duration: 10 s.
40. RateBeautyPic Subjects looked at a photo, and rated how beautiful they thought it was. Duration: 6 s.
41. RateBeautySound Subjects listened to a piece of music, and rated how beautiful they thought it was. Duration: 10 s.
42. RateBeautyMov Subjects viewed a movie, and rated how beautiful they thought it was. Duration: 10 s.
43. RateDisgustPic Subjects looked at a photo, and rated how disgusting they thought it was. Duration: 6 s.
44. RateDisgustSound Subjects listened to a sound, and rated how disgusting they thought it was. Duration: 6 s.
45. RateDisgustMov Subjects viewed a movie, and rated how disgusting they thought it was. Duration: 10 s.
46. RateHappyPic Subjects looked at a photo, and rated how happy the situation seemed to be. Duration: 6 s.
47. RateHappyMov Subjects viewed a movie, and rated how happy the situation seemed to be. Duration: 10 s.
48. RateDeliciousPic Subjects saw a photo of food, and rated how delicious it looked. Duration: 6 s.
49. RateDeliciousMov Subjects viewed a movie of food, and rated how delicious it looked. Duration: 10 s.
50. RatePainfulPic Subjects looked at a photo, and rated how painful the situation seemed to be. Duration: 6 s.
51. RatePainfulMov Subjects viewed a movie, and rated how painful the situation seemed to be. Duration: 10 s.
52. RateNoisy Subjects listened to a sound, and rated how noisy they thought it was. Duration: 8 s.
53. RatePoem Subjects read a poem, and rated how good they thought it was. Duration: 12 s.
54. WordMeaning Subjects judged whether the meaning of a presented word matched the sentence displayed above the word. Duration: 6 s.
55. EyeMoveEasy Subjects looked at a small circle moving around in 1 Hz. Duration: 8 s.
56. EyeMoveHard Subjects looked at a small circle moving around at 2 Hz. Duration: 8 s.
57. WorldName Subjects looked at the photo of a foreign celebrity and judged whether their name matched the name displayed above the photo. Duration: 6 s.
58. DomesticName Subjects looked at the photo of a local celebrity and judged whether their name matched the name displayed above the photo. Duration: 6 s.
59. SoundPlace Subjects listened to an environmental sound, and judged whether it matched the location on the screen. Duration: 6 s.
60. WorldPlace Subjects looked at a photo of a place in some foreign country, and judged whether it matched the site displayed above the photo. Duration: 6 s.
61. DomesticPlace Subjects looked at a photo of a place in their home country, and judged whether it matched the site displayed above the photo. Duration: 6 s.
62. MusicCategory Subjects judged whether the genre of a piece of music matched the name displayed on the screen. Duration: 10 s.
63. DetectTargetPic Subjects judged whether a target item was shown in a photo. Duration: 8 s.
64. DetectTargetMov Subjects judged whether a target item was shown in a movie clip. Duration: 10 s.
65. Metaphor Subjects read a metaphorical text and judged whether the writer’s intention matched the meaning indicated above the text. Duration: 8 s.
66. Irony Subjects read an ironical text and judged whether the writer’s intention matched the meaning indicated above the text. Duration: 8 s.
67. TimeMov Subjects judged whether the duration of a presented movie matched the duration indicated on the screen. Duration: 8 s.
68. TimeSound Subjects judged whether the duration of a presented sound matched the duration indicated on the screen. Duration: 8 s.
69. ComparePeople Subjects looked at two photos of people and judged whether or not the two were the same person. Duration: 6 s.
70. DetectDifference Subjects looked at two pictures and judged whether or not they were exactly the same. Duration: 8 s.
71. Harmony Subjects listened to a sequence of chords and judged whether the chord progression was consonant or dissonant. Duration: 6 s.
72. DecideFood Subjects looked at four photos of different foods and judged which looked the most delicious. Duration: 8 s.
73. DecidePeople Subjects looked at four photos of different people and judged who looked the most reliable. Duration: 8 s.
74. DecidePresent Subjects chose one among four items they wanted to receive as a present. Duration: 8 s.
75. DecideShopping Subjects chose one among four items they would buy during shopping. Duration: 8 s.
76. LanguageSound Subjects listened to a sound and judged whether the language matched the language indicated on the screen. Duration: 6 s.
77. DetectColor Subject judged whether the colour of a word matched the colour displayed above the word. Duration: 6 s.
78. SoundLeft Subjects judged whether a sound was presented from their left side. Duration: 8 s.
79. SoundRight Subjects judged whether a sound was presented from their right side. Duration: 8 s.
80. RelationLogic Subjects read a syllogism based on spatial relationships and indicated whether the conclusion was valid or not. Duration: 12 s.
81. PropLogic Subjects read a syllogism based on prepositional logical relationships and indicated whether the conclusion was valid or not. Duration: 12 s.
82. MoralPersonal Subjects read a text and judged whether the described activity (which included harming somebody) was ethically permissible or not. Duration: 12 s.
83. MoralImpersonal Subjects read a text and judged whether the described activity (which did not include harming somebody) was ethically permissible or not. Duration: 12 s.
84. Recipe Subjects judged whether a given recipe matched the actual recipe of a given dish. Duration: 8 s.
85. TimeValue Subjects selected one of two money rewards which would be offered to them at different points in the future. Duration: 8 s.
86. PressOrdEasy Subjects pressed buttons based on a series of numbers presented at 1 Hz. Duration: 8 s.
87. PressOrdHard Subjects pressed buttons based on a series of numbers presented at 2 Hz. Duration: 8 s.
88. Rhythm Subjects listened to a series of sound pulses and judged whether its rhythm was constant or not. Duration: 6 s.
89. RecallTaskEasy Subjects judged whether the two earlier tasks matched the task described on the screen. Duration: 6 s.
90. RecallTaskHard Subjects judged whether the three earlier tasks matched the task described on the screen. Duration: 6 s.
91. MemoryDigit Subjects memorized a series of digits. Duration: 6 s.
92. MatchDigit Subjects judged whether a presented series of digits matched the one presented before (corresponding to the digits memorized in the MemoryDigit task). Duration: 6 s.
93. MemoryLetter Subjects memorized a series of letters. Duration: 6 s.
94. MatchLetter Subjects judged whether a presented series of letters matched the one presented before (corresponding to the letters memorized in the MemoryLetter task). Duration: 6 s.
95. MemoryNameEasy Subjects memorized three names associated with three photos of different animal species. Duration: 6 s.
96. MatchNameEasy Subjects judged whether two presented photos matched the names displayed on the screen (corresponding to the names memorized in the MemoryNameEasy task). Duration: 8 s.
97. MemoryNameHard Subjects memorized three names associated with three photos of the same animal species. Duration: 6 s.
98. MatchNameHard Subjects judged whether two presented photos matched the names displayed on the screen (corresponding to the names memorized in the MemoryNameHard task). Duration: 8 s.
99. ForeignRead Subjects read an English sentence (i.e. a foreign language for the subjects). Duration: 12 s.
100. ForeignReadQ Subjects answered a question about the English sentence they read just before. Duration: 6 s.
101. ForeignListen Subjects listened to an English sentence. Duration: 10 s.
102. ForeignListenQ Subjects answered a question about the English sentence they listened to just before. Duration: 6 s.
103. MirrorImage Subjects judged whether a photo was symmetrical or not. Duration: 6 s.

